# Fungal and algal lichen symbionts show different transcriptional expression patterns in two climate zones

**DOI:** 10.1101/2025.06.05.657779

**Authors:** Henrique F. Valim, Jürgen Otte, Imke Schmitt

**Affiliations:** Senckenberg Biodiversity and Climate Research Centre, Frankfurt am Main, Germany; LOEWE Centre for Translational Biodiversity Genomics (LOEWE-TBG), Frankfurt am Main, Germany; Goethe University Frankfurt, Institute of Ecology, Evolution and Diversity, Frankfurt am Main, Germany

**Keywords:** light induction, metatranscriptome, gene expression, symbiosis, algae, fungi, Trebouxia, RNA-seq, environmental specialization, mutualism

## Abstract

In the lichen symbiosis, the fungal and algal partners constitute a closely integrated system. The combination of fungal and algal partners changes along climate gradients in many species, and is expected to be adaptive. However, the functional mechanisms behind this symbiosis-mediated environmental adaptation are unknown. We investigate which transcriptional profiles are associated with specific fungal-algal symbiont pairings found in lichens from high elevation (cold temperate) and low elevation (Mediterranean) sites. Using laboratory-acclimatized thalli, we find that lichen fungal and algal symbionts show variable expression profiles between high- and low-elevation individuals: circadian- and temperature-associated genes for fungi and light-responsive genes for algae show site- specific patterns. We also find that high- and low-elevation individuals differentially express sugar and carbohydrate transporters in both fungal and algal symbionts, pointing to symmetrical and climate-dependent carbohydrate transport mechanisms between the symbionts. A light pulse treatment identified asymmetries between fungal and algal light response, with high- and low-elevation fungal symbionts but only low-elevation algal symbionts responding to the light pulse. We identify light responsive genes in fungal symbionts that may differ from those previously described for other fungi. Together, these results shed light on functional differences between environmentally specialized lichen individuals, made up of niche-specific partner combinations.

## Background

Lichens are evolutionarily stable symbioses composed of a primary fungal partner, the mycobiont, and an algal or cyanobacterial partner, the photobiont (Ahmadjian, 1993; Spribille et al., 2022; Pichler et al., 2023). Typically, lichens are understood as a mutualism wherein the photobiont provides a photosynthesis-derived carbohydrate source to the mycobiont, while the mycobiont provides micronutrients, a better hydration regime, protection against grazing, and environmental protection for the photobiont (reviewed in Spribille et al. 2022). Many lichen mycobionts have wide or near-cosmopolitan distributions (Galloway, 2008) and associate with multiple species-level photobiont lineages along their range, e.g., Lecanora rupicola (Blaha et al., 2006), Evernia mesomorpha (Piercey-Normore, 2006), Thamnolia vermicularis (Nelsen and Gargas, 2009), Cetraria aculeata (Fernández- Mendoza et al., 2011), Tephromela atra (Muggia et al., 2014), Ramalina menziesii (Werth and Sork, 2014), and Umbilicaria pustulata (Rolshausen et al., 2018).

Studies along geographic gradients have identified several environmental factors as drivers of photobiont distribution. In the genus Cladonia, 172 mycobiont species associate with 545 lineages of Asterochloris photobionts across a cosmopolitan range, with both climate and fungal identity driving genetic variation in the photobiont lineages (Pino-Bodas and Stenroos, 2021). A narrower analysis of Cladonia spp. within Europe revealed a small number of environmentally-dependent clusters of photobiont lineages, with photobiont switching in hosts only occurring within each cluster (Škvorová et al., 2022). A study of lichens from Cladonia as well as Stereocaulon and Lepraria spp. across the Canary islands, Madeira, Sicily and Aeolian islands also identified that the photobionts from the local pool of algal species was determined by both host specificity (i.e., the total number of mycobionts each photobiont formed symbioses with) and temperature, with different hosts responding differently to the temperature gradient (Vančurová et al., 2021). Similar patterns were observed for photobionts across these same three lichen genera in Bolivia, with lower photobiont specificity and fungal haplotype diversity in cosmopolitan host species (Kosecka et al., 2021).

Three species of the genus Umbilicaria have been extensively studied for their population turnovers along elevation gradients: U. pustulata and U. phaea, which grow in a continuum from Mediterranean to cold temperate climate zones (Hestmark, 1992; Hestmark, 2004; Rolshausen et al., 2022), and U. hispanica, which occurs in the cold temperate and alpine biomes (Sancho and Crespo, 1989). Part of the observed population differentiation of mycobionts in U. pustulata, U. hispanica and U. phaea can be linked to environmental factors, as evidenced by abrupt, genomic breaks along elevation gradients in all three species that separate populations into high- and low-elevation genetic clusters corresponding to different climate zones (Dal Grande et al., 2017; Rolshausen et al., 2020; Rolshausen et al., 2022). These genotypic changes between climate zones at high- and low- elevation have also been identified at loci of functional significance, such as circadian and temperature-associated genes (Valim et al., 2023), secondary metabolite genes (Singh et al 2022), and a kinase with unknown function (Merges et al. 2023). In addition to the genomic differentiation of the mycobiont, these Umbilicaria species also show a turnover of green algal symbionts (Trebouxia species) along elevation (Dal Grande et al., 2018; Rolshausen et al., 2022).

Although patterns of species distributions for myco- and photobionts are beginning to be better described in lichens, understanding the specific functional consequences of these pairings is in its infancy. The physiological properties of lichen thalli from different environments, and with different partner combinations, can vary; for instance, Trebouxia lineages found in U. pustulata thalli from cold temperate and Mediterranean environments have been shown to display different photosynthetic response curves to thallus water content, with individuals from hot and dry environments reaching maximal photosynthesis at a lower thallus water content than individuals from cold and wet environments (Dal Grande et al., 2017). Furthermore, a study on the physiological properties of thalli with different partner combinations has shown that photosynthetic performance depends on both the identity of the algal symbiont and on species and even genotype identity of the fungal host (Schulz et al., 2022).

The metatranscriptional responses of lichen thalli to environmental stimuli have recently also begun to be dissected. In the marine cyanolichen Lichina pygmaea, different cyanobacterial photobionts are more transcriptionally active depending on whether the thallus is hydrated by seawater at high tide or freshwater in the form of rain at low tide (Chrismas et al., 2024). In the photosymbiodeme lichen Peltigera britannica, where different parts of the thallus associate with either a cyanobacterial photobiont only or a combination of cyanobacterial and green algal photobionts, different thermal stress responses were observed between green algal and cyanobacterial symbionts, and mycobionts were found to differentially express antibiotic and growth-inhibitory genes, which may play a role in how the mycobiont manages photobiont communities depending on the presence of green algae (Almer et al., 2023). The transcriptional responses of Evernia mesomorpha myco- and photobionts to liquid water vs water vapor hydration have also recently demonstrated asymmetries in how the two primary partners in the lichen thallus respond to physiological conditions (Meyer et al., 2024).

Although the above studies have demonstrated how different photobionts can affect the expression profile of the lichen mycobiont and vice versa, it remains unclear how symbiont changes along climate gradients affect gene expression. To explore the role of both lichen mycobiont and photobiont identity on transcriptional profiles at the extremes of a climate gradient, we sampled Umbilicaria pustulata thalli from high- and low-elevation sites on Mt. Limbara, Sardinia, Italy, with the high-elevation site (1250 m) being located in the cold temperate climate zone and the low-elevation site (166 m) in the Mediterranean zone. To exclude environmental effects as much as possible, we performed the experiments in a climate chamber after acclimatizing the lichens over a 72h period. To investigate additional physiological and transcriptional differences between high and low elevation samples, such as responses to specific environmental stimuli, we performed a commonly used experimental design for testing light-responsive and circadian-associated gene expression: after dark adaptation, we applied a 20-minute light pulse, which reveals early light responses in both algal and fungal partners.

We asked the following questions: Which mycobiont and photobiont genes are differentially expressed in thalli from high- and low-elevation sites? Are previously-identified genes with allele frequency differences between high- and low-elevation thalli of U. pustulata, such as circadian clock-associated genes, also differently expressed under acclimatized conditions? How do high- and low-elevation individuals differ in their response to environmental stimuli, such as light?

## Methods

### Sample collection, growth conditions, and light pulse treatment

In order to determine the extent of transcriptional variation for both myco- and photobionts in different climate zones, we collected U. pustulata thalli for RNA-seq analysis at the two most extreme points along the climatic gradient: a low-elevation site at Ponte Diana, adjacent to Coghinas lake (166 m, 40°45’26.3333"N, 9°04’41.9322"E, “Ponte Diana” in Fig. 1A) in the Mediterranean climate zone, and a high-elevation site near the peak of Mt. Limbara (1,250 m, 40°51’24.6341"N, 9°10’06.8779"E, “Mt. Limbara” in Fig. 1A) in the cold temperate climate zone. Thalli were collected from sun-exposed granitic rocks on 29 March 2022, air-dried, and transported to a growth chamber in Frankfurt am Main, Germany.

**Figure 1:**
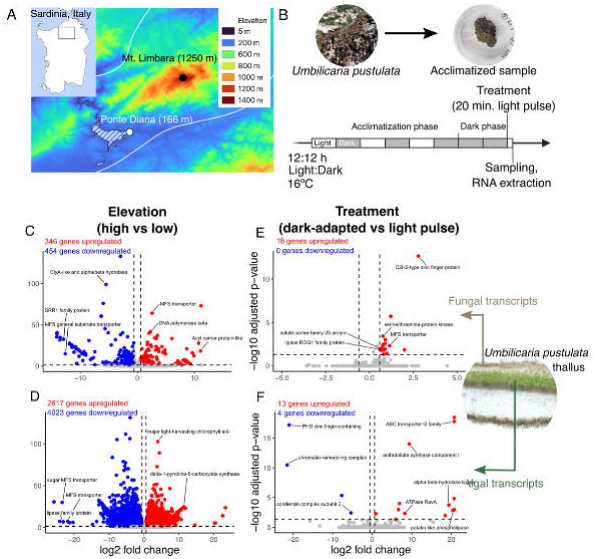
Experimental setup and transcriptional profile of Umbilicaria pustulata samples taken in Sardinia, Italy. (A) Samples were harvested from 2 sites: a high- and low-elevation extremes of a gradient along Mt. Limbara, Sardinia, Italy. (B) Samples were brought back to the lab and acclimatized for 3 days under constant humidity and temperature before a light pulse experiment was performed. Differentially expressed genes identified between high- and low-elevation in fungal (C) and algal (D) transcripts, and between dark-adapted and light pulse-treated samples in fungal (E) and algal (F) transcripts. Horizontal dotted line denotes -log(0.05) and vertical dotted lines denote +/- 1.5 fold change in expression; transcripts beyond these levels are marked red.

After Trebouxia operational taxonomic unit (OTU) identification via Sanger sequencing using a previously-described ITS marker (Kroken and Taylor, 2000; Sadowska- Deś et al., 2013), we selected 4 high-elevation thalli containing the OTU2 Trebouxia lineage (GB accession number MH299133) and 4 low-elevation thalli containing the OTU3 (= “S19” according to the nomenclature in (Muggia et al., 2020) Trebouxia lineage (GB accession number MH299167) for the subsequent experiment. In order to minimize transcriptional variation associated with environmental conditions at each site during sampling, U. pustulata thalli were rehydrated and acclimatized over 72 h in a climate chamber before being subjected to a 32-hour dark adapted phase followed by a 20-minute light pulse treatment and subsequent RNA-seq analysis (Fig. 1B).

Growth conditions and the light pulse treatment were performed as in (Valim et al., 2022). Briefly, dried U. pustulata individuals were separated into two thallus pieces, one for the dark-adapted (DD) and one for the light pulse sample (LL). Thallus pieces were placed in closed 6 cm Petri dishes and rehydrated by applying 0.5 ml dH_2_O to blotting paper and allowing lichen thalli to rehydrate through water vapor, while acclimating to laboratory conditions over 3 days with 12h light/12h dark cycles (60 μmol photons m^-2^ s^-1^) at 16° C. Thalli were kept hydrated with dH_2_O applied to the filter paper each day. After 3 days, samples were then exposed to an additional 24 h of constant darkness, before light pulse-treated samples were exposed to a 20-minute light pulse (60 μmol photons m^-2^ s^-1^) at 8:00 (circadian time 0, subjective dawn) and all samples were harvested immediately into liquid nitrogen and stored at -80° C before RNA extraction.

RNA extraction, library construction and RNA-sequencing After tissue harvesting from lichen thalli (150 mg), RNA was extracted with TRI Reagent (Zymo Research Europe GmbH, Freiburg, DE) according to the manufacturer’s instructions. The extracted RNA was then sent to Novogene (Cambridge, UK) for library construction, quality control and 150 bp paired-end sequencing with NovaSeq. All sequence information can be found on the European Nucleotide Archive (ENA) under project accession number PRJEB72275.

### Differential gene expression analysis

Differential gene expression analysis was done by adapting the pipeline established by Mistry et al. (Mistry et al., 2021). To avoid the quantification of algal reads, the transcripts from the most recently annotated genome of U. pustulata published in (Singh et al., 2022) was used for read mapping. Briefly, after quality control (QC) and trimming of raw reads using FastQC v0.11.9 (Andrews, 2010) and Trimmomatic v0.39 (Bolger et al., 2014), indexing (salmon index) and quantification of RNA transcripts was performed using Salmon v0.13.1 (Patro et al., 2017). QC of the mapped sequences was done using HISAT2 v2.2.1 (Kim et al., 2019) and SAMtools v1.17 (Danecek et al., 2021) to align reads to U. pustulata genome scaffolds, followed by QualiMap 2 v2.3 (Okonechnikov et al., 2016). All QC information was subsequently aggregated and analyzed using MultiQC v1.14. Differential gene expression (DGE) analysis was conducted using DESeq2 v1.38.3 (Love et al., 2014) in R.

Gene IDs were confirmed, when possible, by running protein sequences from genes of interest (e.g., identified by DESeq2 analysis for light treatment and elevation, Tables S1-4) against InterPro protein BLAST, with proteins that could not be identified (%ID below 40%) otherwise labeled as “hypothetical protein.” Conserved domains were identified, when possible, by running protein sequences against NCBI Conserved Domains Search (CDD v3.21).

### Weighted gene co-expression network analysis

The identification of co-expressed genes in parallel to DESeq2 analyses was done using WGCNA (Langfelder & Horvath, 2007) in R. Briefly, after normalization of read counts for all genes, we determined the soft threshold level for the “power” parameter using the pickSoftThreshold function by calculating a measure of the model fit, the signed R^2^, above 0.80. We subsequently ran blockwiseModules (power = 14 for fungal reads, 9 for algal reads; TOMType = “signed”; maxBlockSize = 20000; randomSeed = 1234). We used limma v3.54.2 (Ritchie et al., 2015) in order to determine if any of the identified modules’ eigengenes were significantly correlated to treatment or site.

## Results

### U. pustulata fungal and algal transcriptomic fractions demonstrate elevation-associated and light pulse-responsive transcriptional variation

After mapping to U. pustulata Ascomycota scaffolds derived from metagenomic sequencing (hereafter, fungal transcripts) and Trebouxia sp. scaffolds derived from sequencing pure algal cultures (hereafter, algal transcripts), a total of 8,772 fungal and 17,629 algal transcripts were quantified from the metatranscriptomic dataset. A large number of fungal (800, Fig. 1C and Table S3) and algal (6,640, Fig. 1D and Table S4) transcripts were differentially expressed between high- and low-elevation populations. A small number of fungal (18, Fig. 1E and Table S1) and algal (17, Fig. 1F and Table S2) transcripts were differentially expressed by the 20-minute light pulse (DESeq2 adjusted p-value < 0.05, fold change > 1.5).

Fungal and algal genes show different high- and low-elevation co-expression patterns Previous work has identified large differences in genes linked to carbohydrate and sugar transport among lichen-forming fungi (Resl et al., 2022). Here, we compared the expression levels of sugar, sugar alcohol and carbohydrate transporters in both the fungal and algal fractions of the transcriptome. Sugar transporters were among the most differentially-expressed genes in both the fungal and algal transcripts. We identified a set of 27 sugar and sugar alcohol transporters in the fungal transcripts (Fig. 2A) and a set of 29 carbohydrate and sugar transporters in the algal transcripts (Fig. 2B). For both the fungal and algal transcripts, a sharp division between transporters that were overexpressed in high- vs low-elevation fungal genotypes and algal species were observed, with 17/27 (63%) fungal and 17/29 (59%) algal sugar transporters forming a cluster downregulated in the high-elevation individuals, while 10/17 (37%) fungal and 12/29 (41%) algal sugar transporters forming a cluster downregulated in the low-elevation individuals.

**Figure 2:**
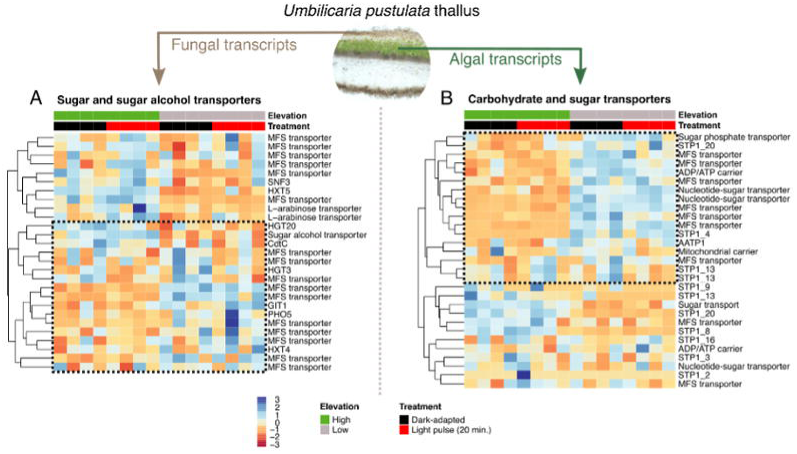
Fungal and algal sugar transport-associated genes are differentially expressed in individuals from high and low elevations. A. fungal reads. B. algal reads. Dashed boxes indicate sugar transporters that are downregulated in high-elevation individuals. A light pulse treatment has no effect on fungal and algal sugar, sugar alcohol and carbohydrate associated gene expression.

Circadian- and temperature-associated genes show elevation-specific co-expression, but not light pulse-dependent patterns We have previously identified a set of 50 circadian- and 37 temperature-associated fungal genes with allelic variation in high- and low-elevation populations of U. pustulata across several elevation gradients, including Mt. Limbara, Sardinia (Valim et al., 2023). Here, we analyzed the expression profile of these gene sets and found a cluster of genes in the circadian-associated (20/52, 38%, Fig. 3A) and temperature-associated (17/37, 46%, Fig. 3C) gene sets that were downregulated in the high-elevation fungal genotype.

**Figure 3:**
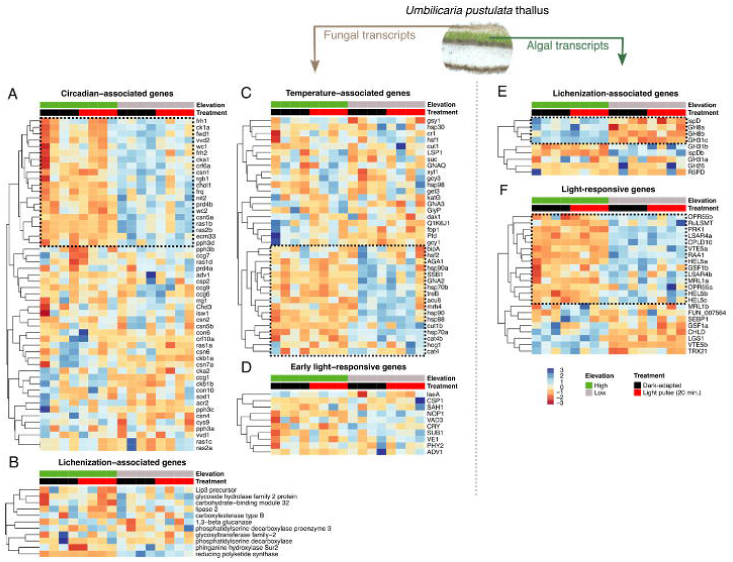
Circadian- and temperature-associated as well as light-responsive genes show different patterns at high- and low-elevation sites for fungal and algal reads. A. Circadian-associated fungal genes identified in Valim et al. (2024). B. Lichenization-associated fungal genes identified in Kono et al. (2020). C. Temperature-associated fungal genes identified in Valim et al. (2024). D. Early light-responsive fungal genes derived from Yu and Fischer (2019). E. Lichenization-associated algal genes identified in Puginier et al. (2024). F. Light-responsive algal genes identified in Fauser et al. (2022). Dashed boxes indicate genes that are downregulated in high-elevation individuals.

We analyzed the expression profile of two additional gene sets of interest, lichenization-associated and light-responsive genes, in both the fungal and algal reads. The algal “lichenization-associated gene set” was derived from a phylogenomic analysis of the evolution of lichenization by (Puginier et al., 2024) and for the fungal reads from a metatranscriptomic study of lichen symbiosis resynthesis experiments by (Kono et al., 2020). The algal light-responsive gene set was derived from homologs of light-sensitive genes characterized in Chlamydomonas reinhardtii (Fauser et al., 2022) and for the fungal reads from well-known early light signaling genes in N. crassa, reviewed in (Yu and Fischer, 2019). Both gene sets demonstrated some algal species-specific clustering: 4/9 of the algal lichenization-associated genes were downregulated in the low-elevation species (Fig. 3E), while 14/22 (64%, Fig. 3F) of the light-responsive genes were downregulated in the high-elevation species. However, neither the lichenization-associated (Fig. 3B) nor the early light responsive fungal genes (Fig. 3D) demonstrated clustering in a genotype- or treatment-specific manner.

Weighted gene co-expression module analysis identifies fungal and algal-specific light pulse-responsive modules In order to further investigate how mycobiont expression patterns were affected by the light treatment or the identity of both the fungal genotype and algal symbiont species, we performed a weighted gene co-expression network analysis (WGCNA) to identify modules of co-expressed genes. We identified 56 modules of co-expressed genes for the fungal transcripts (Fig. S2A-C) and 9 modules for the algal transcripts (Fig. S2D-F). 27/56 (48%) of the fungal gene co-expression modules were significantly correlated (adjusted p value < 0.05) with high- vs low-elevation genotype identity, comprising 3,868/8,772 (44%) of all fungal transcripts quantified (Table S5). 2/9 of the algal gene co-expression modules were significantly correlated (adjusted p value < 0.05) with high- vs low-elevation species identity, comprising 10,384/17,629 (59%) of all algal transcripts quantified (Table S6). To ascertain the overlap between the WGCNA and DESeq2 approaches, we compared transcripts in the largest co-expression modules for both the fungal and algal transcripts with the DESeq2 results for high- vs low-elevation genotype/species. For the fungal genes, 554 transcripts overlapped between the WGCNA module 1 and DESeq2 elevation results, accounting for 45% and 69% of the total genes, respectively (Fig. S3A). For the algal genes, 6,051 transcripts overlapped between the WGCNA module 1 and DESeq2 elevation results, accounting for 66% and 91% of the total genes, respectively (Fig. S4A).

We identified two modules in the fungal transcripts that were significantly differentially expressed by the 20-minute light pulse treatment. Fungal module 18, containing 111 genes (Fig. 4A), was enriched in stress response and transport biological processes (Fig. 4B) as well as a variety of molecular functions associated with the activation of core metabolism, such as oxidoreductase activity, NADPH/NADP^+^ activity and transcription activation (Fig. 4C). Cellular localization results for fungal module 18 point to transcriptional activation at both the nucleus and mitochondrial nucleoid (Fig. 4D). Fungal module 23, containing 84 genes (Fig. 4E), was enriched in a variety of sugar biosynthetic processes and transcriptional activation (Fig. 4F), as well as a variety of molecular functions involved in transcriptional activation and ribosomal activation (Fig. 4G). Cellular localization results for fungal module 23 point to activation of the transcriptional/translational machinery and protein mobilization in the rough endoplasmic reticulum (Fig. 4H).

**Figure 4:**
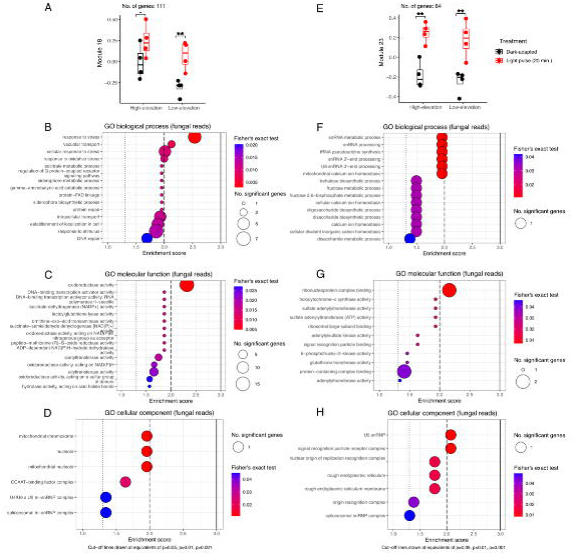
Two co-expression modules in lichen fungal reads characterize early light pulse- dependent responses. A. Plotting of eigengene values for genes in fungal module 18 (111 genes). E. Plotting of eigengene values for genes in fungal module 23 (84 genes). B,F. Gene set enrichment analysis (GSEA) of biological process gene ontology (GO) terms. C,G. GSEA of molecular function GO terms. D,H. GSEA of cellular component GO terms. Dashed and solid lines in GSEA plots denote p =0.05, p=0.01, and p=0.001 for Fisher’s exact test results for each GO term.

We identified one module in the algal transcripts that was significantly downregulated by the 20-minute light pulse in the low-elevation species only. Algal module 6, containing 239 genes (Fig. 5A), was enriched in a variety of ribonucleotide and nucleotide biosynthetic processes (Fig. 5B), transmembrane transporter and nucleotide binding molecular functions (Fig. 5C) that were partly localized to the thylakoids and TRAPPII complex in the protein transport chain (Fig. 5D).

**Figure 5:**
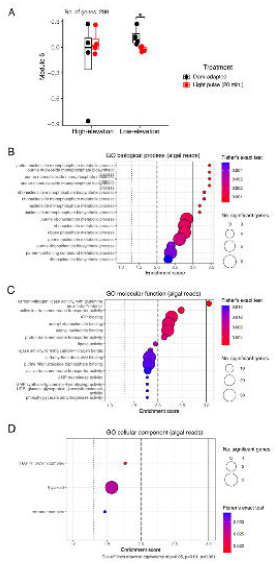
A co-expression module in lichen algal reads reveals species-specific early light pulse-dependent responses. A. Plotting of eigengene values for genes in algal module 6 (239 genes). B. Gene set enrichment analysis (GSEA) of biological process gene ontology (GO) terms. C. GSEA of molecular function GO terms. D. GSEA of cellular component GO terms. Dashed and solid lines in GSEA plots denote p =0.05, p=0.01, and p=0.001 for Fisher’s exact test results for each GO term.

## Discussion

Genotypic variation between high- and low-elevation populations of fungal and algal symbionts dominates expression profiles of U. pustulata.

Previous work in U. pustulata and other Umbilicariacae has painted a clear picture of genotypic variation along elevation gradients in both fungal and algal symbionts (Rolshausen et al., 2020; Valim et al., 2023). However, it has remained unclear whether functional differences exist at the molecular level between lichen individuals from high elevation (with specific symbiont pairings) versus individuals from low elevation (with different, specific symbiont pairings). Here, we report high variation in constitutive gene expression profiles between high- and low-elevation U. pustulata thalli. Constitutive variation in gene expression was much higher between the two algal species, with 6,640 differentially-expressed genes (38% of all algal genes, Fig 1F), compared to only 800 differentially-expressed genes in the two fungal genotypes (9% of all fungal genes, Fig. 1E).

The 800 constitutively DEGs in the fungal symbiont were over-represented in carbohydrate and vitamin/thiamine (vitamin B1) metabolic processes (Fig. S1A), while the 6,640 constitutively DEGs in the algal symbiont were over-represented in a variety of secondary metabolite biosynthesis processes, such as terpenoid and isoprenoid biosynthetic processes (Fig. S1D). Our additional gene co-expression analysis using WGCNA corroborated the DESeq2 pairwise differential expression results, with the two largest modules of co-expressed genes for the fungal symbiont (1,225 genes, Fig. S3A,B) and the algal symbiont (9,149 genes, Fig. S4A,B) each largely overlapping with the results of the DESeq2 analysis.

The overrepresentation analysis for these fungal and algal co-expressed modules identified related but different functional categories: in the fungal co-expressed module, membrane lipid metabolic processes, membrane transporter activity as well as thiamine (vitamin B1) and indole-associated metabolic processes were enriched (Fig. S3C,D), while in the algal co-expressed module, various biosynthetic and metabolic processes and transcriptional activation-associated functional categories were enriched (Fig. S4C,D).

Lichenizing fungi and algae are both known to produce a host of compounds, including phytohormones, that regulate stress responses and may be used for inter-symbiont signaling. Jasmonic acid (JA) and especially Indole-3-acetic acid (IAA), in the auxin class, are known to be produced by lichenizing fungi in culture, with IAA being known to increase Trebouxia water content and fresh mass in biologically-relevant concentrations (Pichler et al., 2020b). Several classic phytohormones, most abundantly IAA but also abscisic acid (ABA), are released by several microalgae, with JA and the gibberellin GA3 also being exuded extracellularly by Trebouxia sp. (Pichler et al., 2020a). These phytohormones are known to mediate fungal (Tsukada et al., 2010) and plant/algal stress responses (Kaur et al., 2022), and may play an additional role in mediating signaling between fungal and algal symbionts. Thiamine/biotin (vitamin B1), which was differentially expressed in the fungal reads, is also known to play a role in oxidative stress responses in fungi (Wolak et al., 2015; Wu et al., 2020). In addition, Major Facilitator Superfamily (MFS) transporters, a ubiquitous class of small-molecule transporters (Pao et al., 1998) that were differentially expressed in both the fungal and algal reads (Fig. 1C-F), may also point to an interplay in small molecules for signaling back and forth between the fungal and algal symbionts.

The classic view of the lichen symbiosis is that the mycobiont benefits by receiving a steady carbon source from the photobiont, both for energy and increased desiccation tolerance, and provides protection from environmental stresses to the photobiont in return (Spribille et al., 2022). Changes in lichen symbiont identity are probably adaptive along climate gradients, but most lichen studies have focused on larger temperate, boreal and alpine lichens in North America and Europe, limiting our knowledge of these dynamics at the physiological and metagenomic level (Stanton et al., 2023). Here, we identify that algal symbionts in particular may be highly variable in their transcriptional capabilities at each extreme of a climate gradient. Previous work has suggested that generalist lichen photobionts are more broadly plastic in their photosynthetic responses (Osyczka and Myśliwa-Kurdziel, 2023). Alongside photobiont identity, this plasticity appears to be strongly influenced by the mycobiont’s protective abilities, e.g., how much arabitol it can provide to photobionts and its effect on light stress under desiccation (Kosugi et al., 2013). Although generalist lichens may be broadly more adapted to a range of conditions than specialized lichens, our findings demonstrate there is likely local adaptation in species with large ranges such as U. pustulata.

U. pustulata light-responsive and circadian clock-associated genes vary between high- and low-elevation sites.

We identified a set of circadian-associated (Fig. 3A) and temperature-associated (Fig. 3C) genes in the fungal symbiont that were constitutively down-regulated in the high-elevation genotype, while a set of light-responsive genes (Fig. 3F) in the algal symbiont was similarly down-regulated in the high-elevation genotype. Curiously, none of these gene sets displayed clear evidence of induction or downregulation by the 20-minute light pulse. The set of fungal circadian genes that was downregulated in the high-elevation genotype included the core circadian clock components frequency (frq), FRQ-interacting RNA helicase (frh), white collar 1 (wc-1) and white collar 2 (wc-2). We have previously analyzed the light pulse activation of frq in U. pustulata (Lecanoromycetes) and the lichenized fungus Dermatocarpon miniatum (Eurotiomycetes); looking at this and the other core circadian clock components more closely, there is a non-significant light pulse effect for frq in the high-elevation genotype, while the low-elevation genotype displays constitutively higher levels of frq expression (Fig. S5A). It may be that the 20-minute light pulse is not sufficient to identify frq light-dependent activation across different fungal genotypes, and that lichen-forming fungi may display slower transcriptional response times than other fungi, like the model species Neurospora crassa (Crosthwaite et al., 1995; Cesbron et al., 2015).

Lichenization-associated and light-sensitive algal genes, but not fungal genes, display strong partitioning between high- and low-elevation thalli.

We analyzed the expression profile of a “lichenization-associated gene set” derived from a phylogenomic analysis of the evolution of lichenization in algae by (Puginier et al., 2024), finding that 4/9 of the lichenization-associated genes were downregulated in the low-elevation species, while the other 5/9 genes were downregulated in the high-elevation species (Fig. 3E). Similarly, 14/22 of the algal light-responsive genes (64%, Fig. 3F) were downregulated in the high-elevation species. By contrast, no clear patterns emerged for the fungal lichenization-associated genes identified by (Kono et al., 2020) (Fig. 3B) or for early fungal light-signaling genes (Fig. 3D).

These stronger elevation-dependent signals for the algal but not the fungal partner may point to differences related to the degree of genetic variation between high- and low-elevation thalli: the fungal partner is highly differentiated along the genome, but is considered the same species, while the algal partners at high and low elevation are considered different species (Rolshausen et al., 2022). In addition, the stronger signals for the algal symbionts may demonstrate variable selection pressures for the algae. This has been hypothesized to occur in lichens, due to the nature of the symbiosis: lichenizing fungi sporulate and recruit local algae at the site of germination when undergoing sexual reproduction, while the algae in the thalli only undergo asexual reproduction and have their growth rates curtailed in symbiosis (Hill, 2009). This latter possibility would be unexpected in

U. pustulata, the lichen under study here, however; this species is known to only undergo asexual reproduction, without sporulation (Hestmark, 1992), which is confirmed by genomic patterns in comparison to a sexually-reproducing species in the same genus, U. phaea (Valim et al., 2023; Wong et al., 2024). More systematic experimental evidence of functional and genomic variation in the algal and fungal fractions of other sexually and exclusively asexually-reproducing lichens along climate gradients, like those of U. phaea and U. pustulata, will need to be gathered to better understand any potential co-evolutionary relationship between lichen symbionts.

Sugar and sugar alcohol transport genes demonstrate climate-specific patterns in both fungal and algal reads.

We also analyzed a variety of gene sets that have been previously identified as being of interest in either lichen research, or fungal/algal research more broadly. Perhaps most interestingly, we observed clear constitutive differences between high- and low-elevation genotypes in fungal and algal sugar transporters, but no activation by the 20-minute light pulse in either (Fig. 2A,B), which was consistent with the results of both the DESeq2 and WGCNA analyses. This is consistent with previous work in E. mesomorpha, which has similarly demonstrated that sugar transporter expression is not drastically affected by other treatments, like shifts from hydration with liquid water versus water vapor (Meyer et al., 2024). We observed that fungal and algal sugar/carbohydrate transporters fall broadly into two co-expressed clusters in the high- or the low-elevation genotypes of fungi and algal species. These results were also consistent with the DESeq2 analysis, where several MFS transporters were identified as among the most differentially-expressed genes in both the fungal and algal fractions of the metatranscriptome (Fig. 1D,E). These genes, part of the Major Facilitator Superfamily, are well-conserved membrane transporters of sugars and sugar alcohols, among other compounds (Pao et al., 1998).

It remains to be studied whether the different sugar transporters being expressed in high- vs low-elevation genotypes of the fungi and algal species are affecting sugar transport dynamics (e.g., different rates of sugar or sugar alcohol transport from algal to fungal partners and vice versa), or whether they might lead to the transport of different sugars or sugar alcohols. Further work should quantify sugar and sugar alcohol concentrations in high- and low-elevation genotypes of U. pustulata under different conditions, both in the field and after acclimatization in the laboratory, to better understand how these may vary along elevation gradients. Fungal-derived sugar alcohols like arabitol have been previously demonstrated to improve desiccation tolerance in lichen-forming algae (Kosugi et al., 2013), which may imply that even among closely related genotypes of lichens living in different climate zones, such as the high- and low-elevation genotypes of U. pustulata at Mt. Limbara, variable sugar transport may be adaptive.

Early light-responsive genes in both fungal and algal symbionts point to a coordinated assimilation response in low- but not high-elevation lichen thalli.

The number of DESeq2-identified genes in the dataset that responded significantly to the brief (20-minute) light pulse was small, with 18 fungal and 17 algal genes (adjusted p-value < 0.05, fold change > 1.5). Of these early light-responsive genes, transporter and lipase-associated biosynthetic genes were identified in both fungal and algal fractions, potentially pointing to a coordinated early response to the light pulse. Interestingly, the WGCNA co-expression analysis identified two additional sets of light-responsive genes in the fungal reads, one with stress response and solute transport-associated genes over-represented (111 genes, Fig. 4A-D) and another with sugar biosynthetic process over-represented (84 genes, Fig. 4E-H). For the algal reads, we identified a single module (239 genes, Fig. 5A-D) of light-responsive genes in the low-elevation thalli only, which was over-represented in transmembrane transporter activity in the thylakoid, perhaps denoting an additional set of early light-activated photosynthesis-associated genes.

Lichen thalli, like other symbioses, are assumed to carefully coordinate responses between symbionts, as has been explored in other symbiotic systems such as the cereal weevil Sitophilus oryzae and its endosymbiont Sodalis pierantonius (Ferrarini et al., 2023). However, recent evidence from lichens points a lack of synchronization in transcriptional responses to different hydration sources in Evernia mesomorpha (Meyer et al., 2024), and the hydration responses of the marine lichen Lichina pygmaea also vary between fresh- and saltwater, with different symbiotic cyanobacteria becoming more transcriptionally active under each hydration regime (Chrismas et al., 2024). Here we observe a similarly asynchronous response to a short light pulse for a high-elevation genotype. Perhaps most interestingly, however, we also observe a more synchronized early light response between fungal and algal symbionts for low-elevation thalli of U. pustulata. Previous work in lichens has identified differential expression profiles and stress responses across different morphs of Peltigera britannica, in which both algal and cyanobacterial symbionts may associate with a single fungal mycobiont (Almer et al., 2023). However, to our knowledge, this is the first evidence of differential responses to a stimulus in lichens containing different species of Trebouxia.

