## Supplemental data for "Fungal and algal lichen symbionts show different transcriptional expression patterns in two climate zones"

**Responses of lichen symbionts to light pulse reveal symmetric expression patterns between fungal and algal genotypes from two climate extremes**

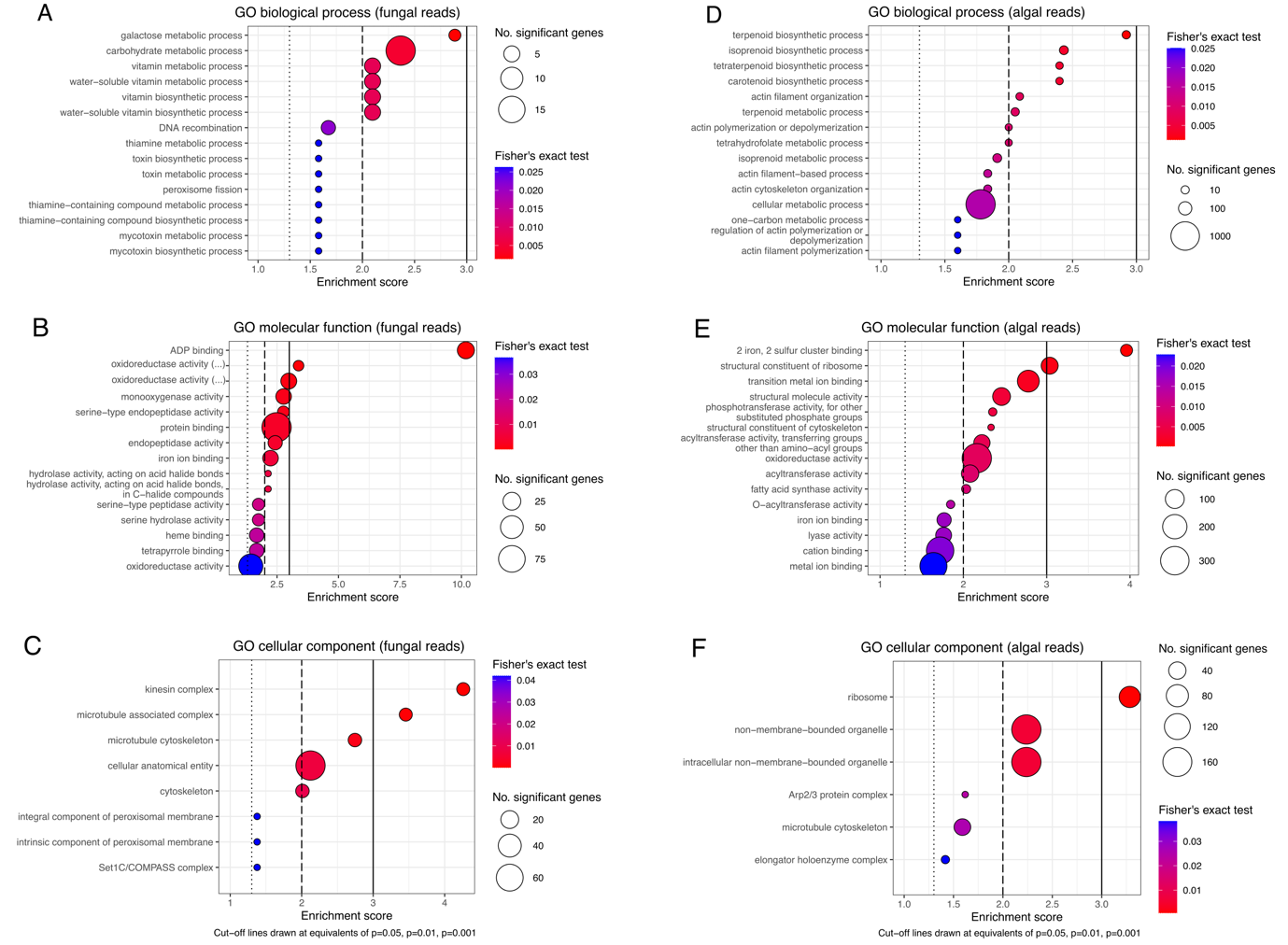

**Figure S1: Gene set enrichement analysis (GSEA) for significant climate-differentiated genes identified in Figure 1 for fungal and algal reads.** A. GSEA of biological process gene ontology (GO) terms for fungal reads. B. GSEA of molecular function GO terms for fungal reads. C. GSEA of cellular component GO terms for fungal reads. D. GSEA of biological process gene ontology (GO) terms for algal reads. E. GSEA of molecular function GO terms for algal reads. F. GSEA of cellular component GO terms for algal reads. Dashed and solid lines in GSEA plots denote p =0.05, p=0.01, and p=0.001 for Fisher’s exact test results for each GO term.

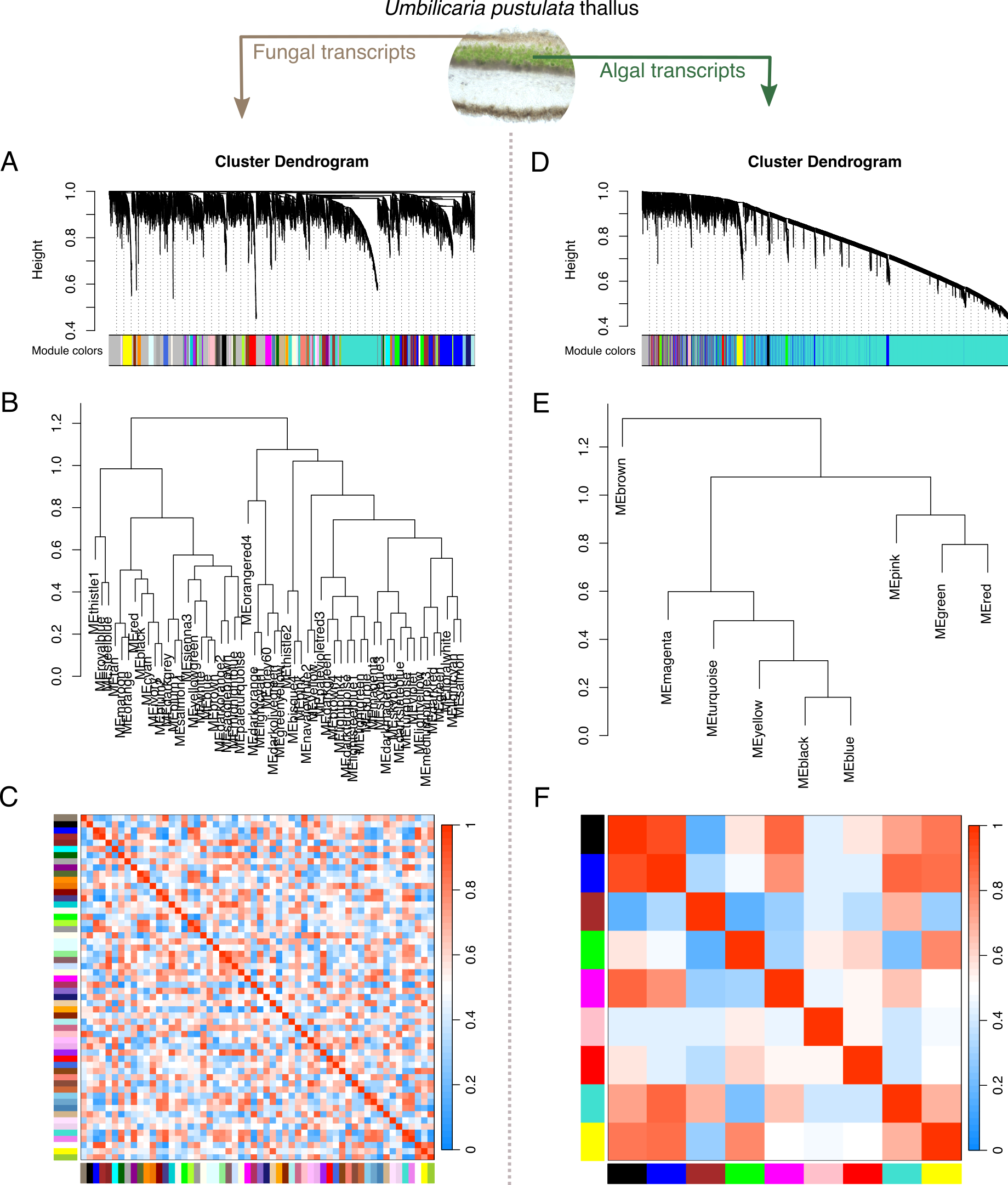

**Figure S2: Results of weighted gene co-expression network analysis (WGCNA) for fungal and algal reads.** Cluster dendogram (A), eigengene dendrogram (B) and eigengene network plot (C) for fungal reads, identifying 56 co-expressed modules. Cluster dendogram (D), eigengene dendrogram (E) and eigengene network plot (F) for algal reads, identifying 9 co-expressed modules.

**Table S1: Fungal transcripts differentially expressed by the 20-minute light pulse (DESeq2 adjusted p-value < 0.05, fold change > 1.5).** Gene: gene ID; Protein classification: protein classification, when available, from InterPro protein BLAST, otherwise labeled as “hypothetical protein”; conserved domains identified: conserved domains identified by NCBI Conserved Domains Search (CDD v3.21), otherwise labeled as “(none)”; baseMean: mean gene expression level of four biological replicates; log2FoldChange: log2 fold change between dark-adapted and 20-minute light treatment induced samples; lfcSE: log2 fold chance standard error; stat: Wald test statistic; pvalue: Wald test statistic p-value; padj: Wald test statistic adjusted p-value.

**Table S2: Algal transcripts differentially expressed by the 20-minute light pulse (DESeq2 adjusted p-value < 0.05, fold change > 1.5).** Gene: gene ID; Protein classification: protein classification, when available, from InterPro protein BLAST, otherwise labeled as “hypothetical protein”; conserved domains identified: conserved domains identified by NCBI Conserved Domains Search (CDD v3.21), otherwise labeled as “(none)”; baseMean: mean gene expression level of four biological replicates; log2FoldChange: log2 fold change between dark-adapted and 20-minute light treatment induced samples; lfcSE: log2 fold chance standard error; stat: Wald test statistic; pvalue: Wald test statistic p-value; padj: Wald test statistic adjusted p-value.

**Table S3: Fungal transcripts differentially expressed between high- and low-elevation (DESeq2 adjusted p-value < 0.05, fold change > 1.5).** Gene: gene ID; Protein classification: protein classification, when available, from InterPro protein BLAST, otherwise labeled as “hypothetical protein”; conserved domains identified: conserved domains identified by NCBI Conserved Domains Search (CDD v3.21), otherwise labeled as “(none)”; baseMean: mean gene expression level of four biological replicates; log2FoldChange: log2 fold change between high- and low-elevation samples; lfcSE: log2 fold chance standard error; stat: Wald test statistic; pvalue: Wald test statistic p-value; padj: Wald test statistic adjusted p-value.

**Table S4: Algal transcripts differentially expressed between high- and low-elevation (DESeq2 adjusted p-value < 0.05, fold change > 1.5).** Gene: gene ID; Protein classification: protein classification, when available, from InterPro protein BLAST, otherwise labeled as “hypothetical protein”; conserved domains identified: conserved domains identified by NCBI Conserved Domains Search (CDD v3.21), otherwise labeled as “(none)”; baseMean: mean gene expression level of four biological replicates; log2FoldChange: log2 fold change between high- and low-elevation samples; lfcSE: log2 fold chance standard error; stat: Wald test statistic; pvalue: Wald test statistic p-value; padj: Wald test statistic adjusted p-value.

**Table S5: WGCNA-identified fungal gene co-expression modules significantly correlated with elevation.** Module: module name; No. genes: number of genes in each module; logFC: log fold change between low- and high-elevation samples. Avg. expression: average normalized expression of all genes in the module. T: Student’s t-test statistic; P value: Student’s t-test p value; adj. P value: adjusted Student’s t-test p value.

| **module** | **No. genes** | **logFC** | **Avg. expression** | **t** | **P value** | **adj. P value** |
| --- | --- | --- | --- | --- | --- | --- |
| ME1 | 1225 | -0.4926561 | 2.95E-17 | -6.2138087 | 8.39E-07 | 4.78E-05 |
| ME36 | 62 | 0.42533875 | 1.10E-16 | 4.64392472 | 6.63E-05 | 0.00188386 |
| ME32 | 67 | -0.4134427 | -4.16E-17 | -4.4253702 | 0.00012178 | 0.00188386 |
| ME52 | 30 | 0.41177537 | -6.94E-18 | 4.39575049 | 0.0001322 | 0.00188386 |
| ME14 | 116 | 0.39370179 | -8.67E-17 | 4.08898173 | 0.00030796 | 0.00275824 |
| ME38 | 56 | 0.39168563 | -1.10E-16 | 4.05627109 | 0.00033681 | 0.00275824 |
| ME3 | 393 | 0.38977319 | 2.39E-18 | 4.02550352 | 0.00036637 | 0.00275824 |
| ME48 | 36 | -0.3885106 | 1.46E-16 | -4.005327 | 0.00038712 | 0.00275824 |
| ME34 | 64 | -0.3839752 | -1.04E-16 | -3.9337267 | 0.00047049 | 0.00282533 |
| ME27 | 73 | 0.38274558 | -6.04E-17 | 3.91454429 | 0.00049567 | 0.00282533 |
| ME49 | 36 | 0.36798688 | 2.38E-16 | 3.69151069 | 0.00090472 | 0.00457907 |
| ME45 | 38 | 0.3663664 | -3.47E-17 | 3.66778907 | 0.00096401 | 0.00457907 |
| ME55 | 23 | 0.35413933 | -5.59E-17 | 3.49329086 | 0.00153238 | 0.00671889 |
| ME53 | 27 | -0.3437055 | 1.13E-17 | -3.3502273 | 0.00222943 | 0.00907696 |
| ME42 | 40 | -0.3356352 | 2.11E-16 | -3.2429484 | 0.00294334 | 0.01118468 |
| ME13 | 122 | -0.3289339 | -1.91E-17 | -3.1559534 | 0.00367858 | 0.01310495 |
| ME37 | 59 | -0.3246836 | -2.30E-17 | -3.1017071 | 0.00422268 | 0.01415839 |
| ME22 | 90 | -0.3205715 | -3.60E-17 | -3.0498842 | 0.00481349 | 0.01524273 |
| ME35 | 62 | 0.31530794 | 3.99E-17 | 2.98446416 | 0.00567199 | 0.01701596 |
| ME43 | 39 | 0.30872896 | -1.07E-16 | 2.90407525 | 0.00692611 | 0.01896353 |
| ME29 | 68 | 0.30843831 | -1.98E-16 | 2.900558 | 0.00698656 | 0.01896353 |
| ME28 | 72 | -0.2978965 | 1.15E-16 | -2.7748622 | 0.00950399 | 0.02462398 |
| ME2 | 686 | 0.29364309 | -2.37E-17 | 2.7251395 | 0.01071727 | 0.02656019 |
| ME31 | 67 | 0.29085488 | 2.69E-17 | 2.69284045 | 0.0115814 | 0.02750582 |
| ME30 | 68 | -0.2861365 | 1.23E-16 | -2.6386999 | 0.01317701 | 0.02977546 |
| ME9 | 134 | -0.2850161 | 2.95E-17 | -2.6259381 | 0.01358179 | 0.02977546 |
| ME15 | 115 | 0.26365006 | 1.21E-16 | 2.38897618 | 0.02352848 | 0.04967124 |
| Sum: | 3868 |  |  |  |  |  |

**Table S6: WGCNA-identified algal gene co-expression modules significantly correlated with elevation.** Module: module name; No. genes: number of genes in each module; logFC: log fold change between low- and high-elevation samples. Avg. expression: average normalized expression of all genes in the module. T: Student’s t-test statistic; P value: Student’s t-test p value; adj. P value: adjusted Student’s t-test p value.

| **module** | **No. genes** | **logFC** | **Avg. expression** | **t** | **P value** | **adj. P value** |
| --- | --- | --- | --- | --- | --- | --- |
| ME1 | 9149 | -0.4994014 | 1.11E-16 | -28.055304 | 1.46E-14 | 1.46E-13 |
| ME2 | 1235 | -0.3668154 | 2.39E-17 | -4.1441926 | 0.00083459 | 0.00417297 |
| Sum: | 10384 |  |  |  |  |  |

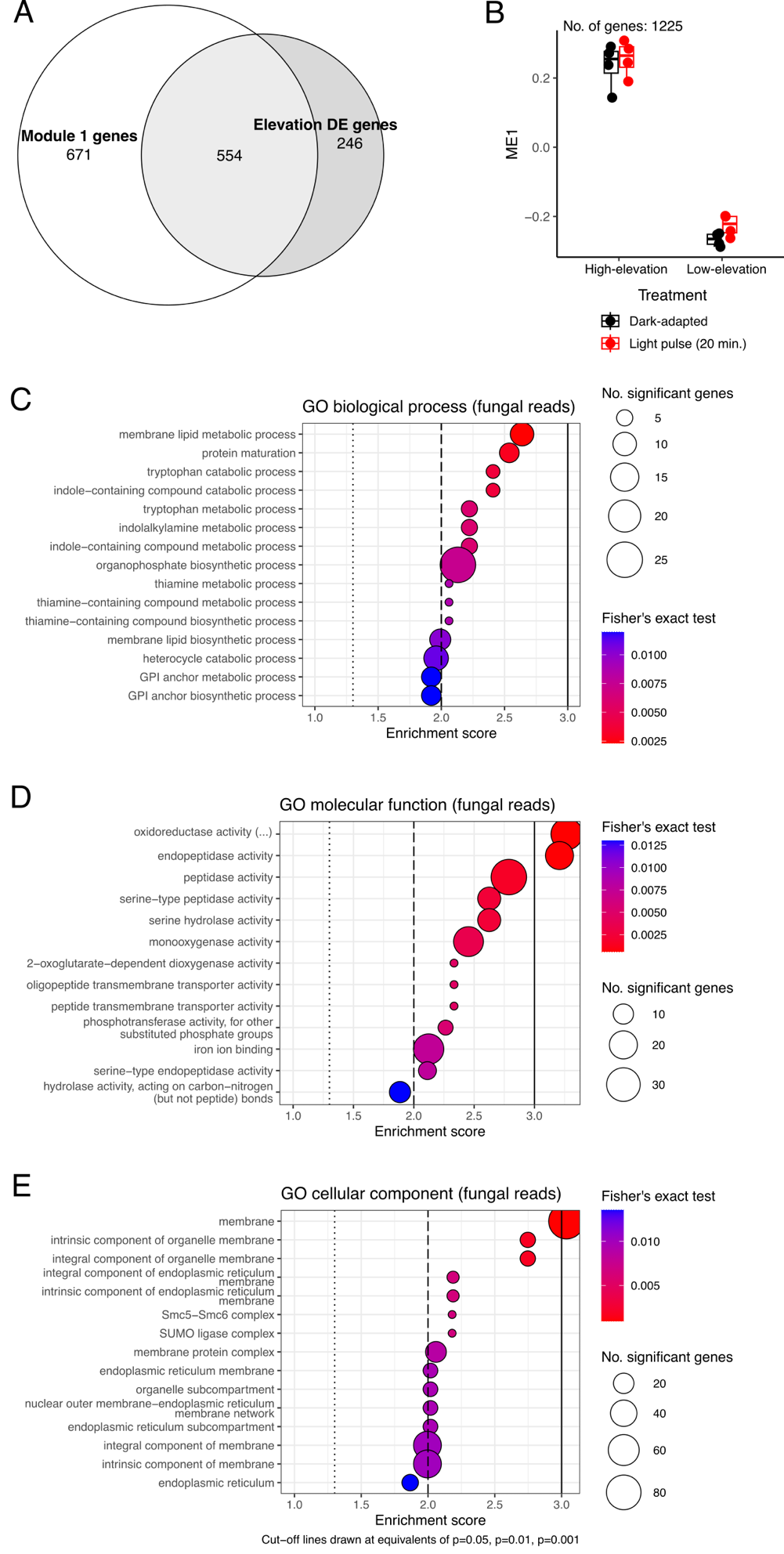

**Figure S3: WGCNA analysis of fungal co-expression module most significantly correlated to elevation (ME1) reveals large overlap with DESeq2 genes that were significantly differentially-expressed (DE) by elevation.** A. Venn diagram of overlap in genes between fungal ME1 and elevation DE genes. B. Plotting of eigengene values for genes in fungal ME1. C. Gene set enrichment analysis (GSEA) of biological process gene ontology (GO) terms. D. GSEA of molecular function GO terms. E. GSEA of cellular component GO terms. Dashed and solid lines in GSEA plots denote p =0.05, p=0.01, and p=0.001 for Fisher’s exact test results for each GO term.

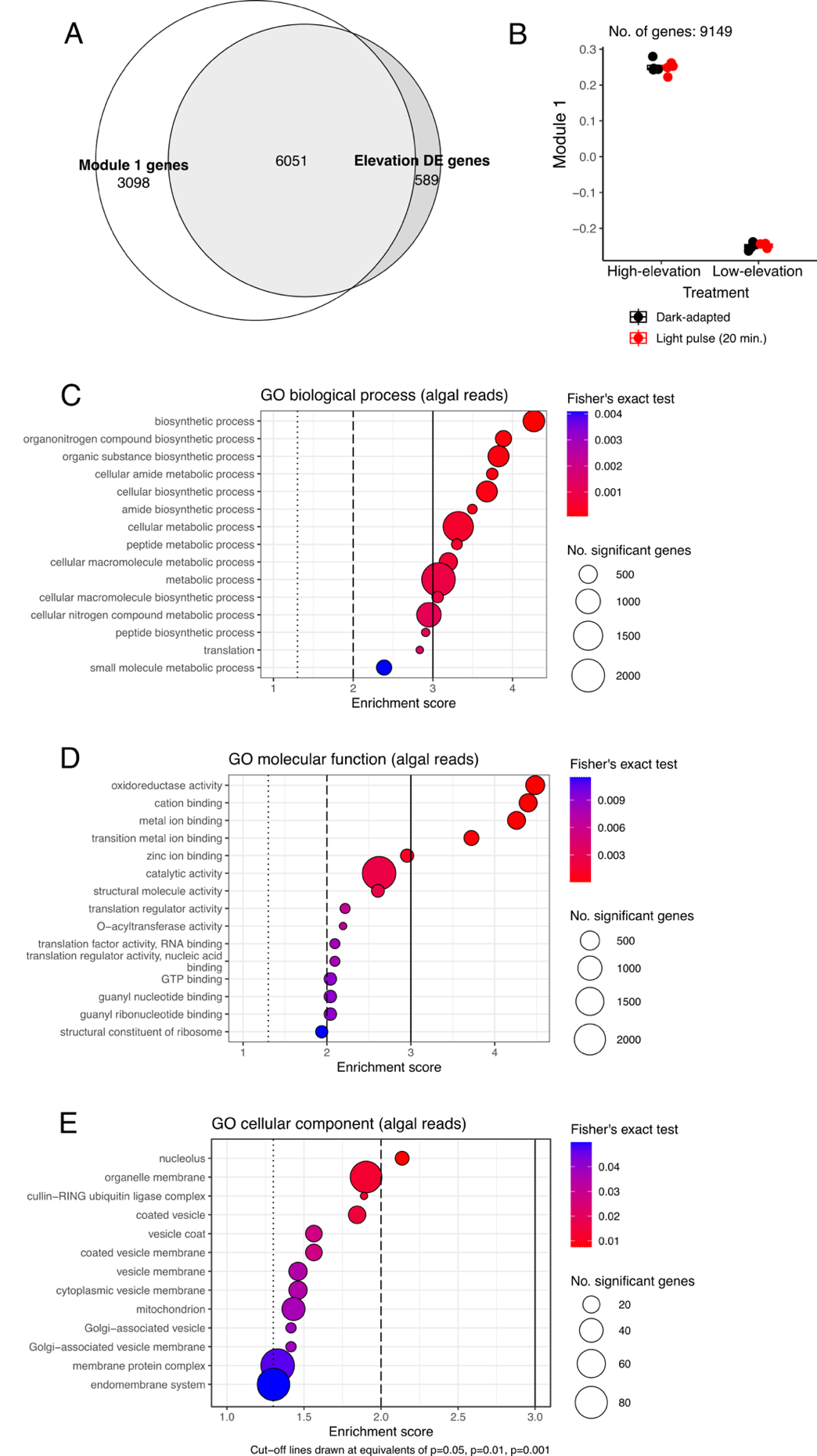

**Figure S4:** **WGCNA analysis of algal co-expression module most significantly correlated to elevation (ME1) reveals large overlap with DESeq2 genes that were significantly differentially-expressed (DE) by elevation.** A. Venn diagram of overlap in genes between algal ME1 and elevation DE genes. B. Plotting of eigengene values for genes in algal ME1. C. Gene set enrichment analysis (GSEA) of biological process gene ontology (GO) terms. D. GSEA of molecular function GO terms. E. GSEA of cellular component GO terms. Dashed and solid lines in GSEA plots denote p =0.05, p=0.01, and p=0.001 for Fisher’s exact test results for each GO term.

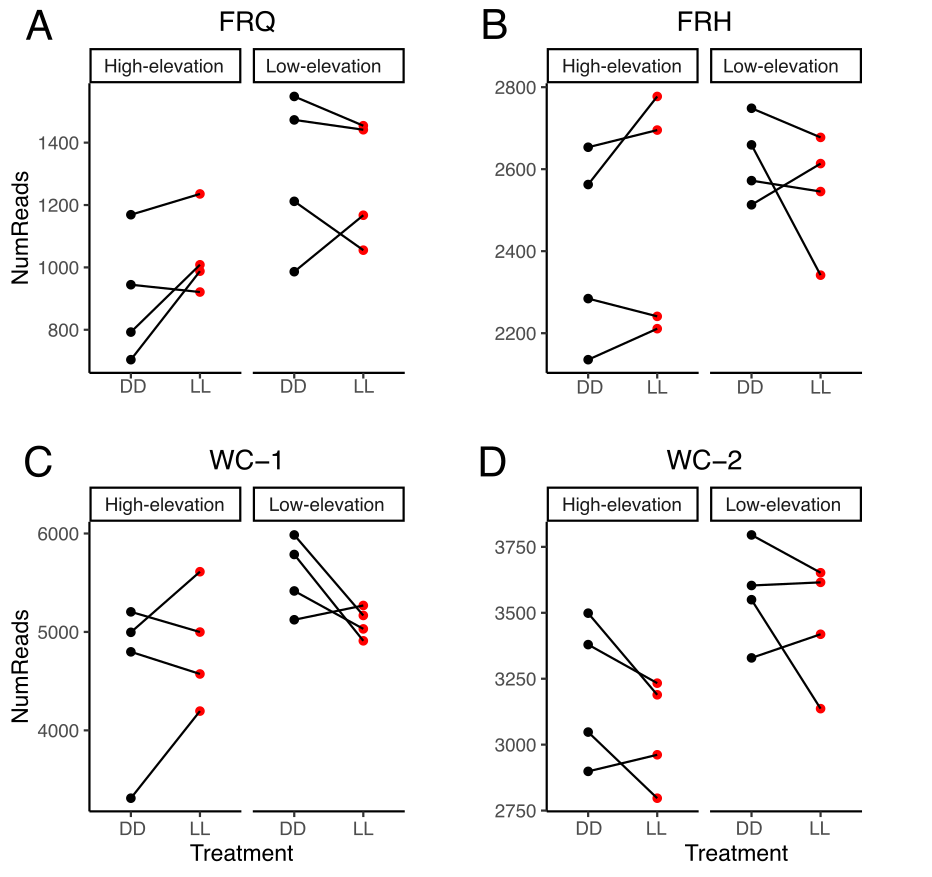

**Figure S5: Paired visualization of core fungal circadian clock gene expression patterns between dark-adapted (DD) and 20-minute light pulse-treated (LL) samples in high- and low-elevation samples.** A. Frequency gene (FRQ). B. Frequency-interacting RNA helicase (FRH). C. White collar 1 (WC-1). D. White collar 2 (WC-2). Solid lines connect same biological individuals between DD and LL treatments.
